# Does Short Molecular Dynamics Rescoring Improve the Ranking of p38*α* Inhibitors over Docking? A Replicate Benchmark

**DOI:** 10.64898/2026.10.27.762277

**Authors:** Akshaya Dhakal, Gungun Chand

## Abstract

Rescoring docked ligands with molecular dynamics (MD) and end point free energy methods such as MM/GBSA is widely used, but its benefit over docking for series of close analogs is uncertain. We compared AutoDock Vina scores with Md based MM/GBSA energies for 29 neutral p38*α* kinase inhibitors (PDB 3FLY) from the Open Force Field protein-ligand benchmark, whose IC50 values from a single publication span 3.8 kcal/mol in estimate ΔG. Each ligand was docked with three random seeds and simulated in two independent 20 ns replicas started from the benchmark pose (AMBER99SB-ILDN, GAFF2, TIP3L; GROMACS). MM/GBSA energies were calculated from 251 frames per trajectory without an entropy term. All 58 simulations remained stable with the ligand bound. Vina scores correlated weakly with experimental ΔG (Spearman *ρ* = 0.28 98% CI -0.15 to 0.62), as did replica averaged MM/GBSA energies (*ρ* = 0.24 -0.18 to 0.62). The difference was -0.03 (-0.50 to 0.43), so the methods were not distinguishable. Independent replicas of the same ligand differed by 3.3 kcal/mol on average, comparable to the experimental range. Raising the interior dielectric constant from 1 to 2 gave a nearly identical ranking (*ρ* = 0.96 between the two sets of scores; *ρ* = 0.37 against experiment, -0.04 to 0.69). In an exploratory analysis, ligands size alone correlated with affinity (*ρ* = 0.60, 0.28 to 0.82), whereas neither computed score tracked size. For this narrow range series, short MD rescoring gave no measurable improvement over docking, and replica variability limited its resolution.

## 1 Introduction

Structure-based virtual screening and lead optimization depend on placing ligands correctly in a binding site and ranking them by affinity. Docking programs such as AutoDock Vina are fast enough for large libraries (1, 2), but their scoring functions are more reliable for pose generation than for affinity ranking (3, 4). The problem is most acute in lead optimization, where close analogs differ by one or two substituents and span a narrow affinity range (5). Resolving differences of a few tenths of a kcal/mol is comparable to both scoring error and experimental uncertainty (6, 7). A common remedy is to rescore docked poses with end-point methods such as MM/GBSA and MM/PBSA, which add force-field energies and continuum solvation at moderate cost (8, 9). Large evaluations show these methods can outper- form standard scoring functions (10), but benefits depend on protocol: the interior dielectric constant matters, and MD-based rescoring may add little over minimized structures (11). MM/GBSA rescoring of docked poses has also been applied to congeneric kinase inhibitor series to predict relative potencies (12). Addressing this requires public, curated structures and affinities for congeneric ligands. The benchmark of Wang et al. (2015) provides this for several targets, including p38α MAP kinase, a well-characterized antiinflammatory target. Most uses of this set concern alchemical free energy calculations (13, 14). Direct comparisons of docking and MD-based MM/GBSA using replicate simulations and explicit uncertainty estimates are rare. With about 30 ligands per target, correlations carry wide confidence intervals, so apparent method differences are easily overstated without resampling statistics. We therefore test whether MM/GBSA rescoring from short MD simulations improves ranking of 29 p38α inhibitors over Vina scores alone. Each ligand was docked with multiple random seeds, and replicate 20 ns simulations were run from benchmark poses. Both methods were compared with experiment by rank and linear correlation with bootstrap confidence intervals and a paired bootstrap for between-method differences. We also assess sensitivity to simulation length and to two ligands near the lower potency limit. All scripts, inputs, and trajectories are provided for reproducibility.

## 2 Methods

### 2.1 Dataset

We used the p38*α* MAP kinase target (PDB 3FLY) from the Open Force Field protein ligand benchmark (15). The 29 neutral ligands have 23-34 heavy atoms and IC50 values of 1-594 nM, all from one publication (16). Protein coordinates and ligand poses were taken from the benchmark files, the poses were generated by the benchmark authors by alignment to the crystallographic ligand, and their metadata notes that some ligands are poorly aligned. Experimental ΔG was estimated as RT ln(IC50) at 298.15 K, an approximation used only for ranking and correlation because IC59 values depend on assay condition.

### 2.2 Docking

The receptor (protein atoms only) and ligands were converted to PDBQT with Open Babel 3.2.1 (17). Ligands were docked with AutoDock Vina 1.2.7 (2) into a 28*20*20 Å box centered at (22.54, 10.62, 31.13) Å, derived from the extent of all benchmark ligand poses plus a 10 Å margin (exhaustiveness 32, 9 modes, seeds 1-3). Scores of the top-ranked mode were averaged over seeds. Pose reproduction was assessed as the symmetry-aware heavy atom RMSD to the benchmark pose in RDkit 2026.03.1 (18), for the crystallographic ligand it was 0.81 Å.

### 2. Molecular dynamics

Each complex (protein with ACE/NME caps, plus ligand), was described with AMBER99SB-ILDN (19), TIP3 water(20), and GAFF2 (21) with AM1-BCC (22)charges generated with Amber- Tools 23.3 (23) via acpype (24). Histidine protonation followed the benchmark structure (HIP at residue 174, HID at 199, 228 and 312, HIE elsewhere), giving a protein charges of -7e (benchmark metadata -5e). System were solvated in a rhombic dodecahedral box (1.0 nm margin)and neutralized with 0.15 M NaCl. After steepest-descent minimization and restrained equilibrium (100 ps NVT at 300K, 200 ps NPT at 1 bar), two independent 20 ns production replicas per ligand were run without restraints in GROMACS 2026.3 (25, 26), starting from the same minimized structure with newly generated velocities (2 fs time step; velocity-rescaling thermostat (27); cell-rescaling barostat (28); LINCS (29); PME (30); 1.0 nm cutoffs; protein and ligand saved every 20 ps). Full parameters are given in Table S4

### 2.4 Trajectory processing

Trajectories were made whole, centered on the protein, and corrected for box jumps. Each had to pass two checks: no protein bond above 2.5 Å(frames sampled every 50th frame) and a protein-ligand center-of-mass distance below 3 nm in all frames (no periodic wrapping). Stability was assessed from backbone RMSD, ligand RMSD after backbone fitting, and minimum protein-ligand distance. All 58 trajectories were retained.

### 2.5 MM/GBSA

Binding energies were computed with gmx_MMPBSA 1.7.0 (31) (MMPBSA.py (32), AmberTools 23.3) from the complex simulation, using 251 frames from 5 to 20 ns. We used the OBC II generalized Born model (igb = 5) (33), 0.150 M salt, interior dielectric 1, the default exterior dielectric, and LCPO surface areas (34) (surface tension 0.0072 kcalmol^*™*1^ Å^*™*2^). ΔTOTAL excludes conformational entropy and is an effective binding energy, not a free energy. A sensitivity analysis used interior dielectric 2. Ligand scores are the mean of two replicas, which were also analyzed separately.

### 2.6 Statstics

Agreement with experimental ΔG was measured by Spearman (primary) and Pearson correlation, with 95% bootstrap confidence intervals (5000 resamples of ligands) and paired bootstrap for the Vina-MM/GBSA difference (10000 resamples). Replica agreement was summarized by mean absolute difference and cross-ligand correlation. The analysis plan (primary outcome: replica averaged MM/GBSA versus Vina; secondary: excluding the two 1 nM ligands, each replica alone; dielectric sensitivity) was written to a dated file after the pipeline had been tested on one ligand and the Vina correlation computed, and before any MM/GBSA score was compared with experiment.

The heavy-atom-count correlation was exploratory. Analyses used SciPy 1.17.1, pandas 3.0.6 and NumPy 2.4.6.

## 3 Results

### 3.1 Dataset

The 29 p38*α* ligands were neutral and contained 23 to 34 heavy atoms. Their IC50 values, all from one publication ranged from 1 to 594 nM, corresponding to estimate ΔG values of -12.3 to -8.5 kcal/mol (range 3.8 kcal/mol). Two ligands (2ee and 21)were at the lowest value, 1 nM (Supplementary table S1).

### 3.2 Docking

Redocking of the crystallographic ligand reproduced its input pose with a heavy atom RMSD of 0.81 Å. For all 29 ligands, the top-scoring Vina pose was within 2 Å of the benchmark pose for 24, 23, and 25 ligands (seeds 1 to 3; median RMSD 0.87–0.90 Å), and for 23 of 29 ligands in all three seeds (Figure S1, Table S3). The largest deviations were observed for 2ee (mean 2.76 Å), 2j (2.53 Å), 2gg (2.34 Å), 2s (2.29 Å), and 2i (2.14 Å). Seeds disagreed noticeably only for 2s (1.70–2.59 Å) and 2n (1.41–2.45 Å). Vina scores were reproducible across seeds (mean range 0.04 kcal/mol, maximum 0.26 kcal/mol) but spanned only 1.9 kcal/mol (*™*10.30 to *™*8.38), half the experimental range.

### 3.3 Stability of the Simulation

All 58 trajectories (29 ligands, two replicas, 20 ns each) remained intact with the ligand bound. The mean protein backbone RMSD per run ranged from 0.15 to 0.29 nm (median 0.18 nm), and the mean ligand RMSD, after fitting on the backbone, from 0.11 to 0.34 nm (median 0.20). The closest protein-ligand atom distance did not exceeded 0.25 nm in any run (Supplementary Table S2). Three runs had a mean ligand RMSD of 0.30 nm or more (2bb, 2i and 2x, replica 2) (Supplementary Figure S2). For 2bb and 2x, the mean ligand RMSD in replica 1 was 0.19 nm.

### 3.4 Correlation with experiment

Results are given in Table 1 and Figures 2 and 3. Vina scores correlated weakly with experimental Δ*G* (Spearman *ρ* = 0.28, 95% CI: *™* 0.15 to 0.62; Pearson *r* = 0.26, 95% CI: *™*0.08 to 0.59). Replica-averaged MM/GBSA energies gave *ρ* = 0.24 (95% CI: *™*0.18 to 0.62; *r* = *™*0.24, 95% CI: 0.20 to 0.60). The difference between the two Spearman coefficients was *™*0.03 (95% CI: *™*0.50 to 0.43), with MM/GBSA higher in 46% of bootstrap resamples; the methods were therefore not distinguishable. Excluding the two 1 nM ligands gave *ρ* = 0.14 (95% CI: *™*0.28 to 0.53) for Vina and *ρ* = 0.27 (95% CI: *™*0.17 to .63) for MM/GBSA. Analyzed separately, replica 1 gave *ρ* = 0.18 (95% CI: *™*0.24 to 0.56) and replica 2 gave *ρ* = 0.24 (95% CI: *™*0.17 to 0.62), so averaging the two replicas did not improve the correlation. All intervals for both methods included zero.

**Table 1.** Correlation of computed scores with experimental Δ*G* (from *IC*_50_) for 29 p38*α* ligands. Spearman *ρ* and Pearson *r* are shown with 95% bootstrap confidence intervals (5,000 resamples of ligands). Scores and Δ*G* are signed so that a positive value means agreement. MM/GBSA energies are Δ*E*_TOTAL_ (251 frames from 5–20 ns, without entropy); “mean of 2 replicas” averages the two independent runs of each ligand. Ligand size is the negative heavy-atom count. Primary: the comparison fixed in advance; secondary and sensitivity: pre-specified; exploratory: not used for any conclusion about the methods.

| Analysis | Score | *N* | Spearman $\rho$ (95% CI) | Pearson $r$ (95% CI) |
| --- | --- | --- | --- | --- |
| Primary | Vina (mean of 3 seeds) | 29 | 0.28 (−0.15, 0.62) | 0.26 (−0.08, 0.59) |
| Primary | MM/GBSA, dielectric 1 (mean of 2 replicas) | 29 | 0.24 (−0.18, 0.62) | 0.24 (−0.20, 0.60) |
| Secondary | MM/GBSA, dielectric 1, replica 1 | 29 | 0.18 (−0.24, 0.56) | 0.20 (−0.22, 0.55) |
| Secondary | MM/GBSA, dielectric 1, replica 2 | 29 | 0.24 (−0.17, 0.62) | 0.22 (−0.21, 0.59) |
| Secondary | Vina, excluding 1 nM ligands | 27 | 0.14 (−0.28, 0.53) | 0.14 (−0.22, 0.52) |
| Secondary | MM/GBSA, dielectric 1, excluding 1 nM ligands | 27 | 0.27 (−0.17, 0.63) | 0.26 (−0.16, 0.60) |
| Sensitivity | MM/GBSA, dielectric 2 (mean of 2 replicas) | 29 | 0.37 (−0.04, 0.69) | 0.32 (−0.11, 0.67) |
| Sensitivity | MM/GBSA, dielectric 2, replica 1 | 29 | 0.30 (−0.12, 0.64) | 0.27 (−0.15, 0.59) |
| Sensitivity | MM/GBSA, dielectric 2, replica 2 | 29 | 0.30 (−0.10, 0.65) | 0.26 (−0.11, 0.61) |
| Exploratory | Ligand size (negative heavy-atom count) | 29 | 0.60 (0.28, 0.82) | 0.54 (0.30, 0.75) |

**Fig 1.**
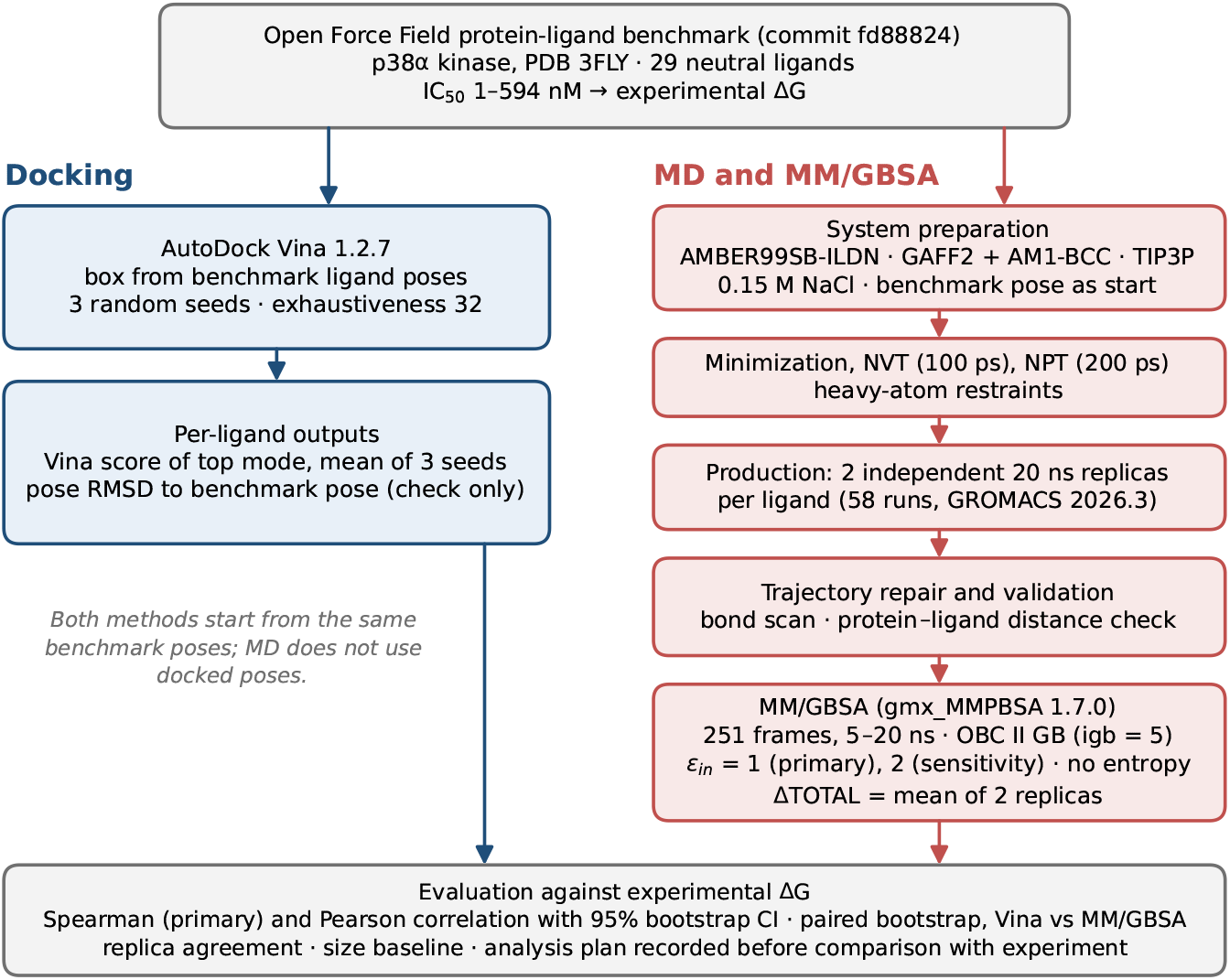
Study workflow. Ligands, structure and IC50 values for 29 p38α inhibitors were taken from the Open Force Field protein-ligand benchmark. Each ligand was docked with AutoDock Vina (three random seeds) and simulated in two independent 20 ns replicas started from the benchmark pose, followed by MM/GBSA rescoring (251 frames, interior dielectric 1; dielectric 2 as a sensitivity analysis). Vina scores and replica-averaged MM/GBSA energies were compared with experimental ΔG by Spearman and Pearson correlation with bootstrap confidence intervals.

**Fig 2.**
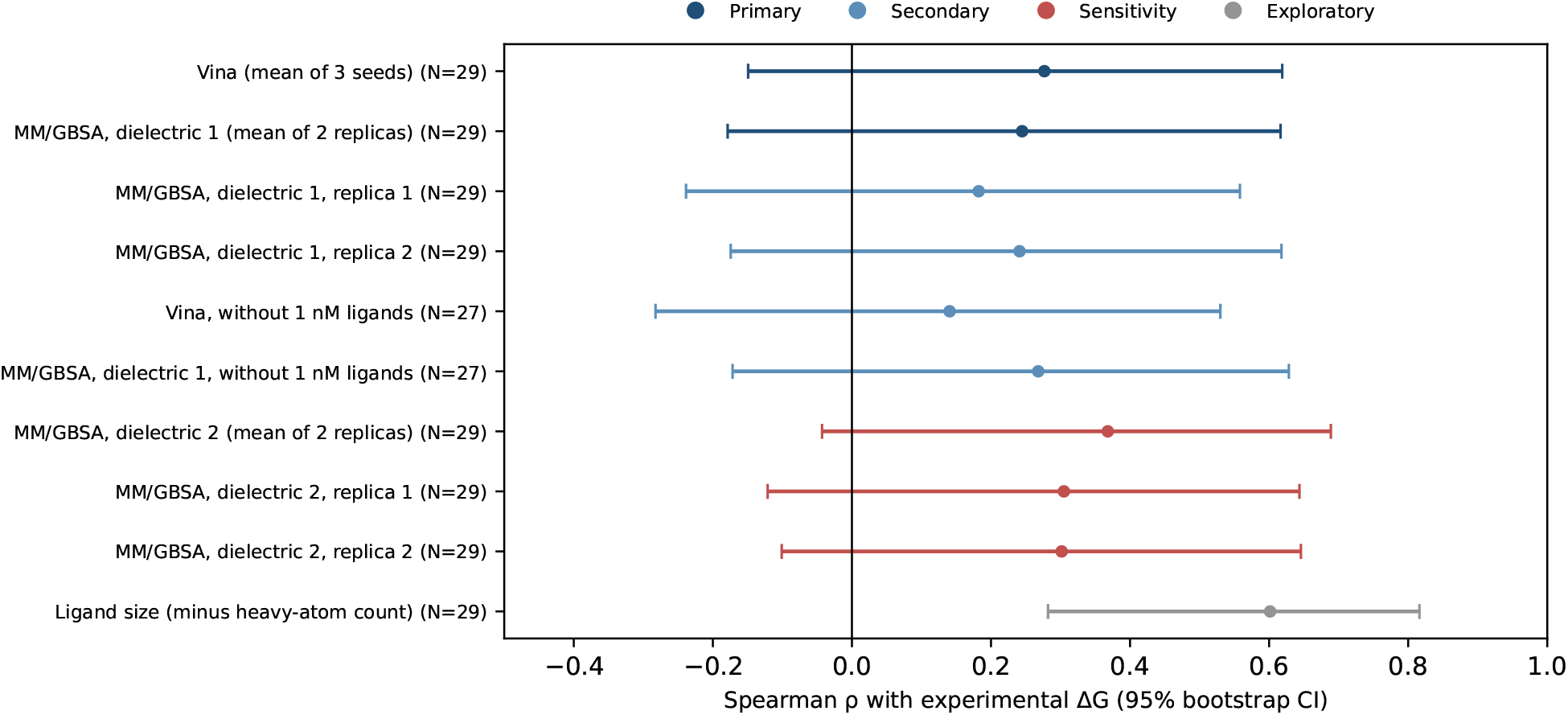
Spearman correlation with experimental ΔG (points) and 95% bootstrap intervals (bars) for each score in Table 1. The vertical line marks zero.

**Fig 3.**
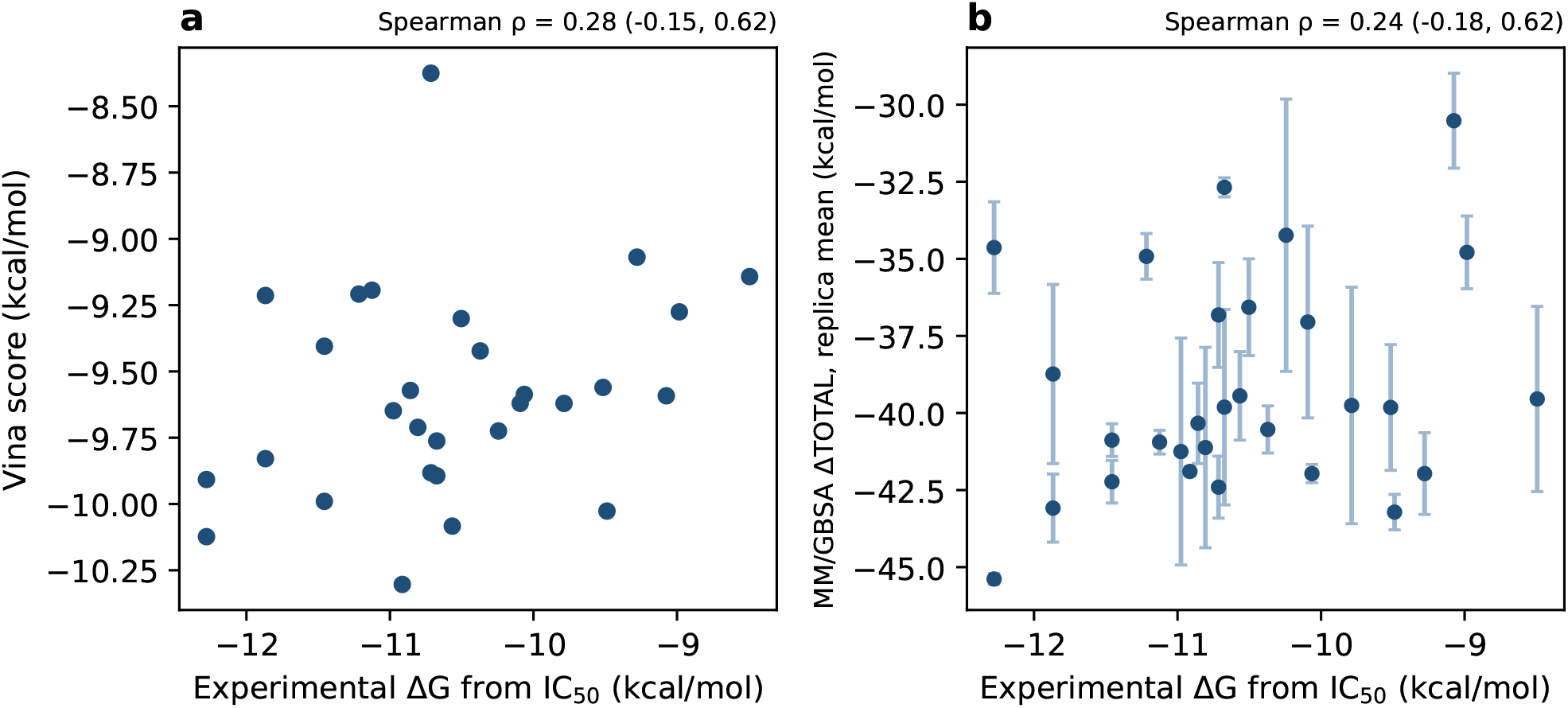
(a) Vina scores and (b) replica-averaged MM/GBSA ΔTOTAL against experimental ΔG. Error bars in (b) show half the difference between the two replicas. Spearman ρ (95% CI) is given above each panel.

### 3.5 Agreement between replicas

The two replicas of the same ligand differed by 3.3 kcal/mol experimental range (Figure 4). Eight ligands differed by more than 5 kcal/mol (2x, 2y, 2m, 2r, 2g, 2j, 2f and 2n) and nine agreed within 1.5 kcal/mol. Replica values correlated only moderately across ligands (Pearson 0.49, Spearman 0.46). For one ligand examined in detail (3FLY), the block standard error within a run was 0.4 kcal/mol, much smaller than the differences between independent replicas.

**Fig 4.**
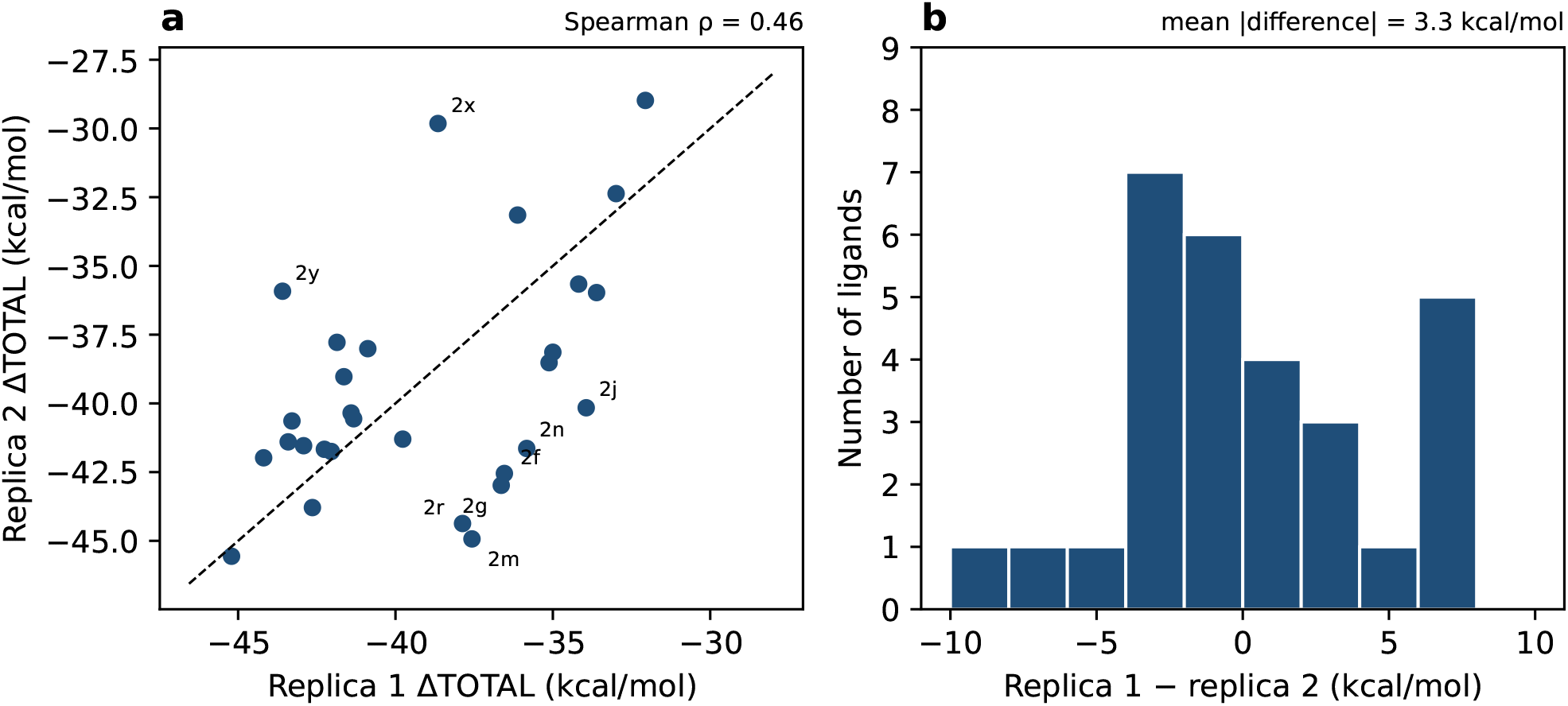
Agreement between replicas. (a) ΔTOTAL of replica 2 against replica 1 (dashed line: identity; ligands differing by more than 5 kcal/mol are labelled). (b) Distribution of the differences between replicas.

### 3.6 Interior dielectric constant and ligand size

Raising the interior dielectric constant from 1 to 2 shifted the mean ΔTOTAL from -39.2 to -45.1 kcal/mol and gave *ρ* = 0.37 (0.04 to 0.69), or 0.30 for each replica alone (Table 1). The replica difference was 3.7 kcal/mol and the ranking was very similar to that at dielectric 1 (*ρ* = 0.96 between the two sets of scores). Experimental ΔG correlated with heavy atom count (*ρ* = *™* 0.60, 95% CI -0.81 to -0.28) (Figure 5), so that ranking ligands by size alone gave *ρ* = 0.60 (0.28 to 0.82)against experiment. Neither Vina scores nor MM/GBSA energies correlated with ligand size (*ρ* = 0.03 for both).

**Fig 5.**
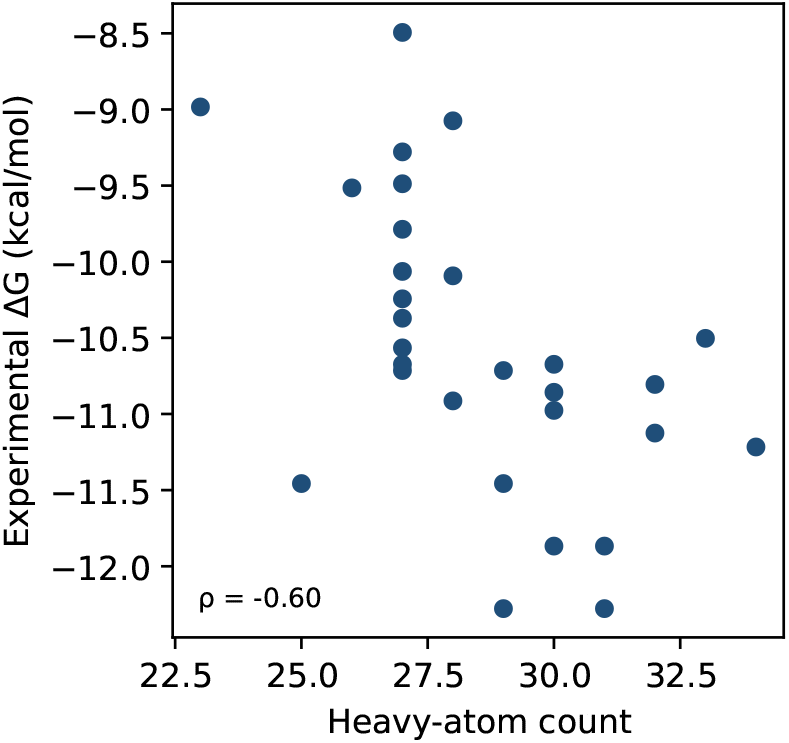
Experimental ΔG against ligand heavy-atom count.

## 4 Discussion

Across a series of 29 closely replated p38*α* inhibitors spanning 3.8 kcal/mol, neither Autodock Vina nor MM/GBSA rescoring from two 20 ns replicas ranked the ligands reliably, and the two methods could not be distinguished. We do not conclude that either method is poor in general. On this set, with this protocol, both correlated weakly and positively with experiment, with intervals reaching zero, an 29 ligand are too few to separate them, the interval for their difference (-0.50 to 0.43) allows a moderate advantage for either.

The best supported explanation is the ratio of signal to noise, which we measured directly. Independent MM/GBSA replicas disagreed by 3.3 kcal/mol on average, as much as the experimental spread, and Vina scores covered only about half of it. No ranking built on values with this variability itself Other factors contributes to this difficulty but cannot be sepated with our data: IC50 values only approximate binding free energies and carry experimental uncertainty (6, 7) the two most potent ligands may sit at the lower limit of assay; MM/GBSA as run here omits conformational entropy and treats the solvent as a continuum, which has been identifies as a source of both under and over estimated affinities (8, 9, 35) and docking scores are known to resolve small substituent changes poorly (3–5)

We consider the variation between replicas the most practically useful result, because it bounds the accuracy of this protocol. Single-trajectory MD-based binding energies have been reported to differ by many kcal/mol between repeats of the same system, which has motivated ensemble approaches that average independent replicas differing only in initial velocities (36–38). Our two replica protocol is consistent with that picture. In one large-scale study of ensemble protocols the error more than halved between ensembles of 5 and 25 replicas (37), and for a DNA-intercalator system the reproducibility of end-point energies depended more on the number of replicas than on the length of each run (39). These observations come from other systems and methods and may not transfer to p38α, but they suggest that more short replicas, rather than longer single runs, would be the more promising way to reduce the variability we observed. Whether our variability reflects insufficient sampling or true ligand mobility remains unresolved: for two ligands (2bb and 2x) the replica with the larger ligand RMSD also gave a less favorable energy, consistent with slow rearrangement of the ligand, but several ligands with large replica differences (such as 2y and 2m) showed no such sign. The block standard errors within single runs greatly understated the variation between replicas, so within run error estimates alone should not be used to judge the reliability of such calculations.

Poor starting poses do not explain the weak correlations. Most Vina poses agreed with the benchmark poses, and all simulations started from the benchmark poses regardless of the docking result. However, only the 3FLY ligand has a crystallographic pose, the other are modeled poses that benchmark’s authors flags as imperfectly aligned in places, so the comparison measures agreement with a model. The optimal interior dielectric constant for MM/GBSA has been reported to vary between systems, with values from 1 to 4 performing best depending on the data set (10, 11, 40), which is why we included it as a sensitivity analysis. The numerically higher correlation at a dielectric of 2 (ρ = 0.37) is in the direction reported for docking rescoring (11) but should not be read as an improvement here. Both MM/GBSA variants were computed from the same trajectories and gave almost the same ranking (ρ = 0.96 between them), the confidence intervals overlap, replica agreement did not improve, and a dielectric of 1, the program default, was fixed as the primary setting before any comparison with experiment. The sensitivity analysis shows that our conclusion does not depend on this parameter. Ligand size correlated with affinity more strongly than either computed score, and the interval for this baseline, unlike theirs, excluded zero. This is plausible in a series where potency follows the size of added substituents, and earlier work has shown that the apparent correlation of scoring functions with experiment often tracks the correlation of ligand molecular weight with affinity (41). It suggests that benchmarks on analog series should report a simple size baseline alongside the correlation of a scoring method. This was an exploratory analysis, and a difference in whether intervals include zero is not a test of a difference between predictors, so we do not claim that size is superior to either method. Previous evaluations reported that MM/GBSA can outperform docking scoring functions on sets of 98 and 855 diverse complexes (10, 35), and it has been applied to congeneric kinase inhibitor series (12), while MD sampling has been reported to add little over minimized structures (11). A narrow series of analogs is a more demanding test than a diverse set, and our results are compatible with each of these findings. We examined only one MM/GBSA protocol apart from the dielectric constant. In particular, MM/PBSA has been described as better suited than MM/GBSA to ranking ligands within a single target (11), and we did not test it. Al- chemical free energy calculations, which were designed for this kind of congeneric ranking and are the main use of this benchmark (5, 14), were not part of our comparison.

## 5 Conclusions

For this p39*α* series, 20 ns MM/GBSA rescoring showed no measurable advantage over docking scores. Independent replicas disagreed by about as much as the experimental affinities span, ligand size alone ranked the compounds at least as well as either method. These results argue for caution when applying short MD rescoring to narrow range analog series, and for reporting replicate variability and a simple size baseline in benchmarks of this kind.

## 6 Code and Data Availability

The scripts used for molecular docking, molecular dynamics analysis, MM/GBSA calculations, statistical analysis, and figure generation, together with the relevant configuration files and processes analysis outputs are available at the GitHub repository: https://github.com/Akshayadhakal/p38_mmgbsa_benchmark. The repository includes documentation describing the computational workflow and software environments. The availability of third-party data and structures is subject to their original sources and applicable reuse conditions.

## Supplementary Material

**Table S1.** Ligand data: experimental IC50 and Δ*G*_exp_ (kcal/mol), heavy atom count (HA), Vina docking score (kcal/mol), Vina pose RMSD (Å), and MM/GBSA binding free energies (kcal/mol) at *ε*_in_ = 1 and *ε*_in_ = 2.

| Ligand | $IC_{50}$ | Unit | $\Delta G_{exp}$ | HA | Vina | Pose RMSD | MM/GBSA ($\epsilon_{in} = 1$) | | | MM/GBSA ($\epsilon_{in} = 2$) | | |
| --- | --- | --- | --- | --- | --- | --- | --- | --- | --- | --- | --- | --- |
|  |  |  |  |  |  |  | Rep 1 | Rep 2 | Mean | Rep 1 | Rep 2 | Mean |
| 2ee | 1 | nM | -12.28 | 31 | -9.91 | 2.76 | -36.12 | -33.15 | -34.64 | -41.55 | -39.72 | -40.64 |
| 2l | 1 | nM | -12.28 | 29 | -10.12 | 0.63 | -45.21 | -45.56 | -45.38 | -51.68 | -52.03 | -51.86 |
| 2n | 2 | nM | -11.87 | 31 | -9.21 | 1.76 | -35.83 | -41.64 | -38.74 | -41.92 | -48.14 | -45.03 |
| 2t | 2 | nM | -11.87 | 30 | -9.83 | 1.00 | -44.19 | -41.98 | -43.08 | -50.68 | -47.65 | -49.16 |
| 3fly | 4 | nM | -11.46 | 25 | -9.40 | 0.81 | -41.41 | -40.35 | -40.88 | -46.54 | -45.93 | -46.24 |
| 2ff | 4 | nM | -11.46 | 29 | -9.99 | 0.78 | -42.92 | -41.54 | -42.23 | -48.77 | -47.19 | -47.98 |
| 2p | 6 | nM | -11.22 | 34 | -9.21 | 1.87 | -34.18 | -35.66 | -34.92 | -41.61 | -43.42 | -42.52 |
| 2s | 7 | nM | -11.13 | 32 | -9.19 | 2.29 | -41.33 | -40.56 | -40.94 | -48.40 | -47.59 | -48.00 |
| 2m | 9 | nM | -10.98 | 30 | -9.65 | 0.64 | -37.57 | -44.93 | -41.25 | -42.69 | -51.85 | -47.27 |
| 3fln | 10 | nM | -10.91 | 28 | -10.30 | 0.35 | -42.04 | -41.75 | -41.90 | -47.83 | -47.71 | -47.77 |
| 3fmh | 11 | nM | -10.86 | 30 | -9.57 | 1.65 | -41.64 | -39.03 | -40.34 | -48.19 | -44.77 | -46.48 |
| 2r | 12 | nM | -10.81 | 32 | -9.71 | 1.47 | -37.87 | -44.37 | -41.12 | -43.91 | -51.21 | -47.56 |
| 3flw | 14 | nM | -10.71 | 29 | -8.38 | 0.85 | -43.41 | -41.40 | -42.40 | -49.99 | -47.30 | -48.64 |
| 2e | 14 | nM | -10.71 | 27 | -9.88 | 0.35 | -35.12 | -38.52 | -36.82 | -39.69 | -42.89 | -41.29 |
| 2gg | 15 | nM | -10.67 | 30 | -9.76 | 2.34 | -32.99 | -32.37 | -32.68 | -38.50 | -38.03 | -38.26 |
| 2g | 15 | nM | -10.67 | 27 | -9.89 | 0.34 | -36.64 | -42.98 | -39.81 | -41.34 | -49.22 | -45.28 |
| 2k | 18 | nM | -10.57 | 27 | -10.08 | 0.33 | -40.88 | -38.01 | -39.44 | -47.30 | -43.71 | -45.50 |
| 2o | 20 | nM | -10.50 | 33 | -9.30 | 1.31 | -35.00 | -38.14 | -36.57 | -43.06 | -46.69 | -44.88 |
| 2z | 25 | nM | -10.37 | 27 | -9.42 | 0.54 | -39.77 | -41.30 | -40.53 | -45.56 | -47.48 | -46.52 |
| 2x | 31 | nM | -10.24 | 27 | -9.72 | 0.45 | -38.65 | -29.82 | -34.24 | -44.78 | -34.34 | -39.56 |
| 2j | 40 | nM | -10.09 | 28 | -9.62 | 2.53 | -33.94 | -40.16 | -37.05 | -39.62 | -45.33 | -42.48 |
| 2i | 42 | nM | -10.06 | 27 | -9.59 | 2.14 | -42.26 | -41.67 | -41.96 | -47.71 | -46.98 | -47.34 |
| 2y | 67 | nM | -9.79 | 27 | -9.62 | 0.89 | -43.59 | -35.92 | -39.76 | -49.15 | -39.86 | -44.50 |
| 3flz | 106 | nM | -9.52 | 26 | -9.56 | 0.50 | -41.86 | -37.78 | -39.82 | -46.88 | -42.51 | -44.70 |
| 2h | 111 | nM | -9.49 | 27 | -10.03 | 1.00 | -42.64 | -43.79 | -43.22 | -48.10 | -49.65 | -48.88 |
| 2aa | 158 | nM | -9.28 | 27 | -9.07 | 0.83 | -43.29 | -40.64 | -41.96 | -47.80 | -45.51 | -46.66 |
| 2bb | 223 | nM | -9.07 | 28 | -9.59 | 1.35 | -32.06 | -28.98 | -30.52 | -39.49 | -36.67 | -38.08 |
| 2v | 260 | nM | -8.98 | 23 | -9.28 | 0.47 | -33.61 | -35.97 | -34.79 | -38.37 | -41.23 | -39.80 |
| 2f | 594 | nM | -8.49 | 27 | -9.14 | 0.99 | -36.54 | -42.55 | -39.54 | -41.11 | -48.73 | -44.92 |

**Table S2.** Summary of MD runs (two replicas per ligand): backbone RMSD, ligand RMSD, and closest protein–ligand distance (nm); mean and maximum over each trajectory.

| Ligand | Replica | Backbone RMSD (nm) |  | Ligand RMSD (nm) |  | Closest distance (nm) |  |
| --- | --- | --- | --- | --- | --- | --- | --- |
|  |  | Mean | Max | Mean | Max | Mean | Max |
| 2aa | 1 | 0.210 | 0.314 | 0.200 | 0.336 | 0.186 | 0.233 |
| 2aa | 2 | 0.184 | 0.280 | 0.188 | 0.345 | 0.187 | 0.241 |
| 2bb | 1 | 0.184 | 0.283 | 0.187 | 0.344 | 0.189 | 0.233 |
| 2bb | 2 | 0.248 | 0.374 | 0.319 | 0.553 | 0.201 | 0.241 |
| 2e | 1 | 0.219 | 0.322 | 0.200 | 0.349 | 0.197 | 0.245 |
| 2e | 2 | 0.233 | 0.367 | 0.212 | 0.446 | 0.196 | 0.235 |
| 2ee | 1 | 0.182 | 0.278 | 0.187 | 0.309 | 0.194 | 0.239 |
| 2ee | 2 | 0.242 | 0.359 | 0.281 | 0.433 | 0.212 | 0.251 |
| 2f | 1 | 0.236 | 0.380 | 0.237 | 0.363 | 0.195 | 0.244 |
| 2f | 2 | 0.165 | 0.259 | 0.201 | 0.313 | 0.201 | 0.240 |
| 2ff | 1 | 0.167 | 0.261 | 0.173 | 0.353 | 0.194 | 0.237 |
| 2ff | 2 | 0.173 | 0.284 | 0.161 | 0.279 | 0.193 | 0.239 |
| 2g | 1 | 0.230 | 0.368 | 0.231 | 0.417 | 0.197 | 0.242 |
| 2g | 2 | 0.157 | 0.205 | 0.147 | 0.235 | 0.198 | 0.232 |
| 2gg | 1 | 0.226 | 0.346 | 0.221 | 0.383 | 0.194 | 0.243 |
| 2gg | 2 | 0.219 | 0.336 | 0.252 | 0.387 | 0.193 | 0.233 |
| 2h | 1 | 0.191 | 0.254 | 0.181 | 0.336 | 0.197 | 0.238 |
| 2h | 2 | 0.168 | 0.234 | 0.167 | 0.248 | 0.195 | 0.234 |
| 2i | 1 | 0.177 | 0.275 | 0.293 | 0.411 | 0.201 | 0.240 |
| 2i | 2 | 0.194 | 0.287 | 0.305 | 0.430 | 0.201 | 0.234 |
| 2j | 1 | 0.205 | 0.301 | 0.221 | 0.369 | 0.199 | 0.238 |
| 2j | 2 | 0.175 | 0.307 | 0.270 | 0.364 | 0.202 | 0.242 |
| 2k | 1 | 0.156 | 0.238 | 0.140 | 0.245 | 0.197 | 0.235 |
| 2k | 2 | 0.192 | 0.287 | 0.182 | 0.323 | 0.196 | 0.237 |
| 2l | 1 | 0.169 | 0.245 | 0.139 | 0.274 | 0.197 | 0.233 |
| 2l | 2 | 0.161 | 0.246 | 0.146 | 0.252 | 0.197 | 0.233 |
| 2m | 1 | 0.250 | 0.357 | 0.244 | 0.386 | 0.193 | 0.234 |
| 2m | 2 | 0.175 | 0.228 | 0.180 | 0.258 | 0.205 | 0.237 |
| 2n | 1 | 0.288 | 0.420 | 0.281 | 0.476 | 0.190 | 0.236 |
| 2n | 2 | 0.163 | 0.235 | 0.188 | 0.278 | 0.196 | 0.232 |
| 2o | 1 | 0.248 | 0.343 | 0.233 | 0.406 | 0.200 | 0.238 |
| 2o | 2 | 0.186 | 0.255 | 0.184 | 0.290 | 0.201 | 0.241 |
| 2p | 1 | 0.234 | 0.340 | 0.283 | 0.418 | 0.196 | 0.239 |
| 2p | 2 | 0.197 | 0.358 | 0.252 | 0.470 | 0.199 | 0.233 |
| 2r | 1 | 0.162 | 0.241 | 0.227 | 0.391 | 0.196 | 0.233 |
| 2r | 2 | 0.161 | 0.250 | 0.211 | 0.278 | 0.195 | 0.232 |
| 2s | 1 | 0.179 | 0.265 | 0.242 | 0.420 | 0.193 | 0.233 |
| 2s | 2 | 0.154 | 0.224 | 0.179 | 0.265 | 0.194 | 0.244 |
| 2t | 1 | 0.183 | 0.257 | 0.111 | 0.242 | 0.197 | 0.240 |
| 2t | 2 | 0.192 | 0.341 | 0.165 | 0.341 | 0.195 | 0.237 |
| 2v | 1 | 0.185 | 0.277 | 0.178 | 0.292 | 0.188 | 0.240 |
| 2v | 2 | 0.181 | 0.283 | 0.142 | 0.269 | 0.192 | 0.238 |
| 2x | 1 | 0.162 | 0.246 | 0.193 | 0.304 | 0.194 | 0.234 |
| 2x | 2 | 0.271 | 0.396 | 0.344 | 0.518 | 0.196 | 0.237 |
| 2y | 1 | 0.174 | 0.232 | 0.237 | 0.306 | 0.188 | 0.225 |
| 2y | 2 | 0.228 | 0.320 | 0.251 | 0.379 | 0.190 | 0.230 |
| 2z | 1 | 0.175 | 0.260 | 0.229 | 0.302 | 0.204 | 0.237 |

Table S2 (continued)
| Ligand | Replica | Backbone RMSD (nm) |  | Ligand RMSD (nm) |  | Closest distance (nm) |  |
| --- | --- | --- | --- | --- | --- | --- | --- |
|  |  | Mean | Max | Mean | Max | Mean | Max |
| 2z | 2 | 0.157 | 0.228 | 0.228 | 0.341 | 0.205 | 0.243 |
| 3fln | 1 | 0.177 | 0.254 | 0.133 | 0.237 | 0.197 | 0.237 |
| 3fln | 2 | 0.163 | 0.256 | 0.121 | 0.230 | 0.197 | 0.235 |
| 3flw | 1 | 0.182 | 0.274 | 0.176 | 0.242 | 0.191 | 0.227 |
| 3flw | 2 | 0.160 | 0.264 | 0.191 | 0.305 | 0.192 | 0.230 |
| 3fly | 1 | 0.192 | 0.280 | 0.174 | 0.284 | 0.200 | 0.237 |
| 3fly | 2 | 0.201 | 0.267 | 0.162 | 0.247 | 0.200 | 0.239 |
| 3flz | 1 | 0.182 | 0.296 | 0.164 | 0.266 | 0.196 | 0.229 |
| 3flz | 2 | 0.196 | 0.314 | 0.239 | 0.446 | 0.193 | 0.231 |
| 3fmh | 1 | 0.154 | 0.247 | 0.210 | 0.340 | 0.196 | 0.229 |
| 3fmh | 2 | 0.177 | 0.292 | 0.201 | 0.344 | 0.194 | 0.236 |

**Table S3.** Docking pose RMSD (Å) relative to the reference pose for three independent docking seeds, with the mean.

| Ligand | Seed 1 (Å) | Seed 2 (Å) | Seed 3 (Å) | Mean (Å) |
| --- | --- | --- | --- | --- |
| 2ee | 2.74 | 2.80 | 2.75 | 2.76 |
| 2j | 2.32 | 2.60 | 2.68 | 2.53 |
| 2gg | 2.32 | 2.34 | 2.37 | 2.34 |
| 2s | 2.59 | 2.58 | 1.70 | 2.29 |
| 2i | 2.14 | 2.14 | 2.13 | 2.14 |
| 2p | 1.87 | 1.86 | 1.87 | 1.87 |
| 2n | 1.41 | 2.45 | 1.41 | 1.76 |
| 3fmh | 1.67 | 1.66 | 1.64 | 1.65 |
| 2r | 1.50 | 1.46 | 1.45 | 1.47 |
| 2bb | 1.35 | 1.33 | 1.35 | 1.35 |
| 2o | 1.31 | 1.31 | 1.32 | 1.31 |
| 2t | 1.00 | 1.00 | 1.00 | 1.00 |
| 2h | 1.00 | 0.98 | 1.01 | 1.00 |
| 2f | 0.98 | 1.00 | 0.98 | 0.99 |
| 2y | 0.90 | 0.90 | 0.87 | 0.89 |
| 3flw | 0.87 | 0.84 | 0.85 | 0.85 |
| 2aa | 0.82 | 0.82 | 0.86 | 0.83 |
| 3fly | 0.81 | 0.81 | 0.82 | 0.81 |
| 2ff | 0.78 | 0.78 | 0.78 | 0.78 |
| 2m | 0.65 | 0.64 | 0.64 | 0.64 |
| 2l | 0.63 | 0.64 | 0.63 | 0.63 |
| 2z | 0.52 | 0.54 | 0.55 | 0.54 |
| 3flz | 0.48 | 0.49 | 0.53 | 0.50 |
| 2v | 0.47 | 0.48 | 0.47 | 0.47 |
| 2x | 0.49 | 0.54 | 0.33 | 0.45 |
| 2e | 0.35 | 0.35 | 0.35 | 0.35 |
| 3fln | 0.35 | 0.35 | 0.35 | 0.35 |
| 2g | 0.33 | 0.33 | 0.34 | 0.34 |
| 2k | 0.32 | 0.33 | 0.34 | 0.33 |

**Fig S1.**
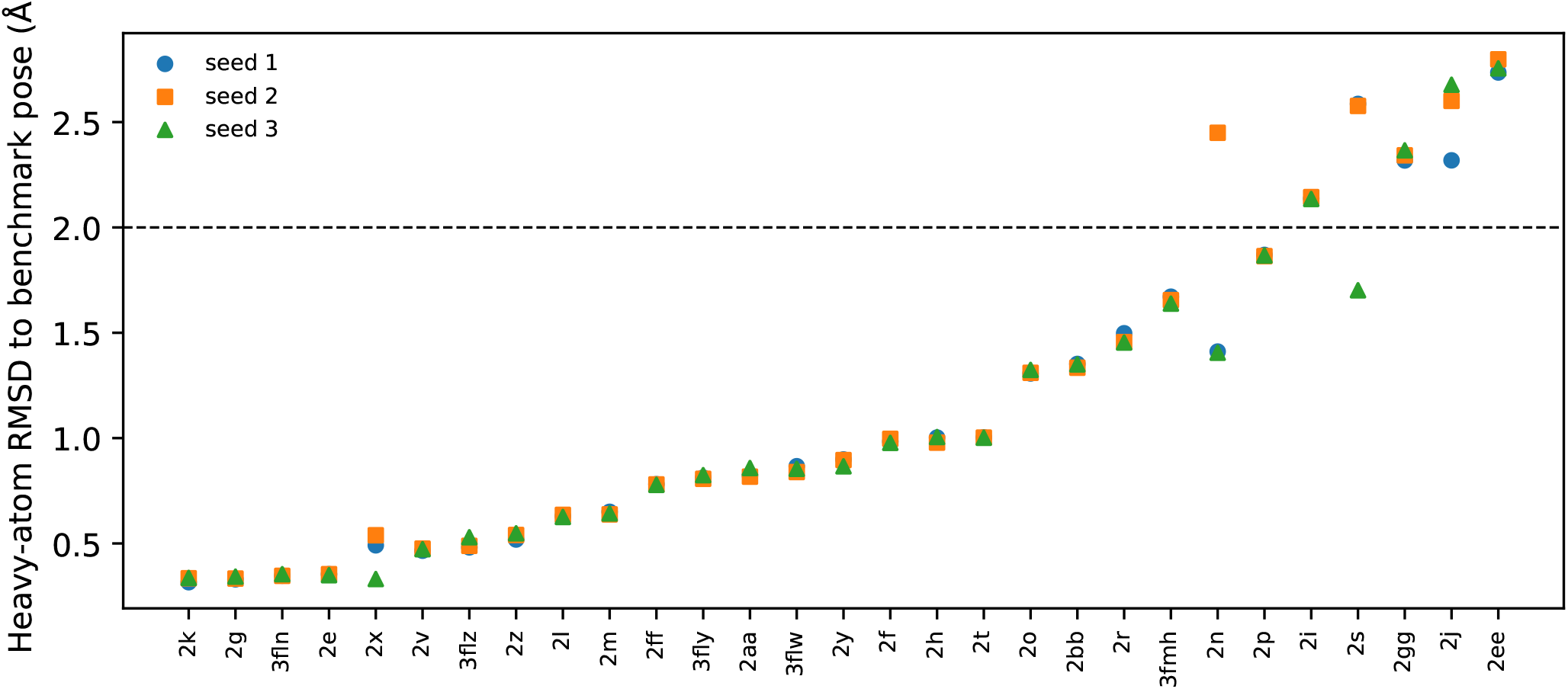
Heavy-atom RMSD of the top Vina pose to the benchmark pose for each ligand and seed; the dashed line marks 2 Å.

**Fig S2.**
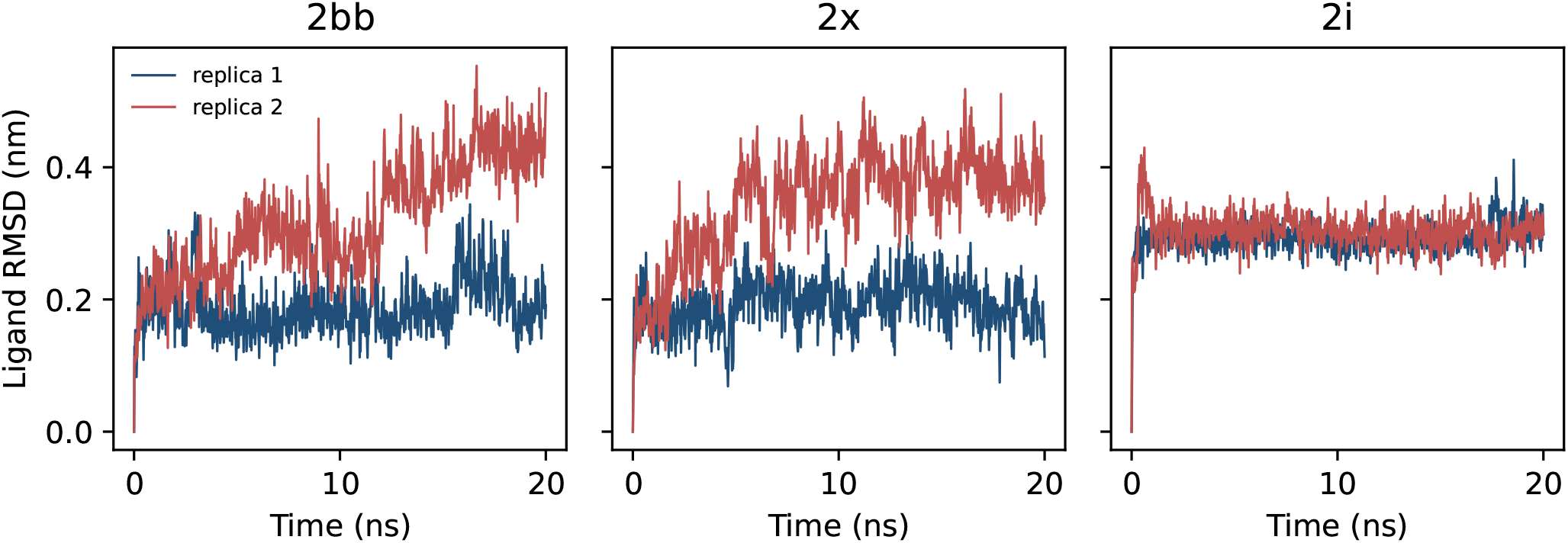
Ligand RMSD (after fitting on the protein backbone) over time for 2bb, 2x and 2i, replicas 1 and 2.

